# A comprehensive analysis of racial disparities in chemical biomarker concentrations in United States women, 1999-2014

**DOI:** 10.1101/746867

**Authors:** Vy Kim Nguyen, Adam Kahana, Julien Heidt, Katelyn Polemi, Jacob Kvasnicka, Olivier J. Jolliet, Justin A. Colacino

## Abstract

**Background:** Stark racial disparities in disease incidence among American women remains a persistent public health challenge. These disparities likely result from complex interactions between genetic, social, lifestyle, and environmental risk factors. The influence of environmental risk factors, such as chemical exposure, however, may be substantial and is poorly understood.

**Objectives:** We quantitatively evaluated chemical-exposure disparities by race/ethnicity and age in United States (US) women by using biomarker data for 143 chemicals from the National Health and Nutrition Examination Survey (NHANES) 1999-2014.

**Methods:** We applied a series of survey-weighted, generalized linear models using data from the entire NHANES women population and age-group stratified subpopulations. The outcome was chemical biomarker concentration and the main predictor was race/ethnicity with adjustment for age, socioeconomic status, smoking habits, and NHANES cycle.

**Results:** The highest disparities across non-Hispanic Black, Mexican American, Other Hispanic, and other race/multiracial women were observed for pesticides and their metabolites, including 2,5-dichlorophenol, o,p’-DDE, beta-hexachlorocyclohexane, and 2,4-dichlorophenol, along with personal care and consumer product compounds. The latter included parabens, monoethyl phthalate, and several metals, such as mercury and arsenic. Moreover, for Mexican American, Other Hispanic, and non-Hispanic black women, there were several exposure disparities that persisted across age groups, such as higher 2,4- and 2,5-dichlorophenol concentrations. Exposure differences for methyl and propyl parabens, however, were the starkest between non-Hispanic black and non-Hispanic white children with average differences exceeding 4 folds.

**Discussions:** We systematically evaluated differences in chemical exposures across women of various race/ethnic groups and across age groups. Our findings could help inform chemical prioritization in designing epidemiological and toxicological studies. In addition, they could help guide public health interventions to reduce environmental and health disparities across populations.

## 1. Introduction

The stark racial disparities in disease incidence and health outcomes among American women remains a persistent public health challenge. For example, preterm birth incidence was found to be approximately 60% higher in non-Hispanic Black women relative to non-Hispanic white women (Culhane and Goldenberg 2011). Non-Hispanic Black and Hispanic women are at increased risk of being diagnosed with developing dysglycemia (Marcinkevage et al. 2013) and diabetes (Cowie et al. 2009), relative to non-Hispanic white women. Non-Hispanic Black women are also 2-3 times more likely to develop the most aggressive subtype of breast cancer, triple negative, compared to non-Hispanic white women (Carey et al. 2006; Stark et al. 2010). Furthermore, relative to non-Hispanic white women, non-Hispanic Black women are also 2.4 times more likely to die of breast cancer after being diagnosed with the pre-invasive lesion, ductal carcinoma *in situ* (Narod et al. 2015).

Recent statistics from the American Cancer Society show variation in trends in breast cancer incidence rates by race/ethnicity in US women from 2005-2014. Specifically, they show increasing trends in breast cancer over time in Asian (1.7% per year), non-Hispanic Black (0.4% per year), and Hispanic (0.3% per year) women, and stable trends in non-Hispanic white and American Indian/Alaska Native women (DeSantis et al. 2017). Dementia rates also vary by race/ethnicity, with rates highest in non-Hispanic black women, followed by American Indian/Alaskan native, Latina, Pacific Islander, non-Hispanic white, and lowest in Asian American women (Mayeda et al. 2016). These rates vary 60% between African American and Asian American women. Reproductive outcomes are also significantly different by race/ethnicity, with studies reporting increased incidence of gestational diabetes in South and Central Asian American women (Thorpe et al. 2005) and Black and Hispanic women (Tanaka et al. 2007). Collectively, these findings suggest profound racial disparities in disease outcomes that manifest throughout the life course. Understanding the etiological factors driving these health disparities is essential for informing public health interventions seeking to promote health equity.

While health disparities are likely due to complex interactions between genetic, social, and lifestyle factors, the impact of genetic factors on disease disparities appears to be minor (Braun 2007; Cooper et al. 2003; Diez Roux 2012). For example, a meta-analysis of genetic factors underlying racial disparities in cardiovascular disease failed to identify heterogeneity of genetic risk factors by race/ethnicity (Kaufman et al. 2015). These findings of a modest genetic impact on differential cardiovascular disease risk by race/ethnicity are consistent with genome-wide association studies. A study found that variants with the strongest association with blood pressure explain, in aggregate, less than 5% of the phenotypic variance (Ehret et al. 2011). Moreover, a meta-analysis of genetic risk factors and cancer disparities reported similar findings, with almost no heterogeneity in cancer risk alleles by race/ethnicity (Jing et al. 2014).

Environmental risk factors may be more influential in generating health disparities than other risk factors. For instance, estimates of environmental impacts on chronic disease suggest than 70-90% of risk is due to environmental exposures (Lim et al. 2012; Rappaport and Smith 2010). A mechanistic understanding of racial disparities in disease therefore requires a characterization of differences in environmental risk factors. In particular, differences in chemical exposures have been hypothesized to be important etiologic factors in racial disparities of disease rates (Hoover et al. 2012; Juarez and Matthews-Juarez 2018; Ruiz et al. 2018; Wang et al. 2016; Zota and Shamasunder 2017).

To investigate the influence of environmental risk factors on health disparities, the goal of this study was to conduct a comprehensive analysis of racial disparities in chemical biomarker concentrations among US women. For this, we leveraged data from the National Health and Nutrition Examination Survey (NHANES), an ongoing population-based health study conducted by the US Centers for Disease Control and Prevention (CDC). Additionally, we developed visuals to highlight differences in biomarker concentrations across races and age groups, by defining the relative magnitude of exposure disparities for individual chemicals and chemical families.

## 2. Methods

### 2.1 Study Population

NHANES is a cross-sectional study designed for collecting data on demographic, socioeconomic, dietary, and health-related characteristics in the non-institutionalized, civilian US population. For this analysis, we used the continuous NHANES data on chemical biomarkers and demographics, which were collected from 1999-2014 with 82,091 participants initially. We excluded participants for not having any data on chemical biomarkers (*N* = 7,001), resulting in a sample size of 75,090 study participants. Since this analysis is focused on measuring chemical disparities in US women, we excluded male participants (*N* = 37,010), leading to a final sample size of 38,080 female participants. For a given chemical, we also excluded participants with missing data on any of the following covariates: race/ethnicity, age, NHANES cycles, poverty income ratio, cotinine levels, and urinary creatinine. Number of participants with complete data for a given chemical and the listed covariates are tabulated in **Excel Table S1**. These exclusion and inclusion criteria are delineated in **Figure 1**.

**Figure 1.**
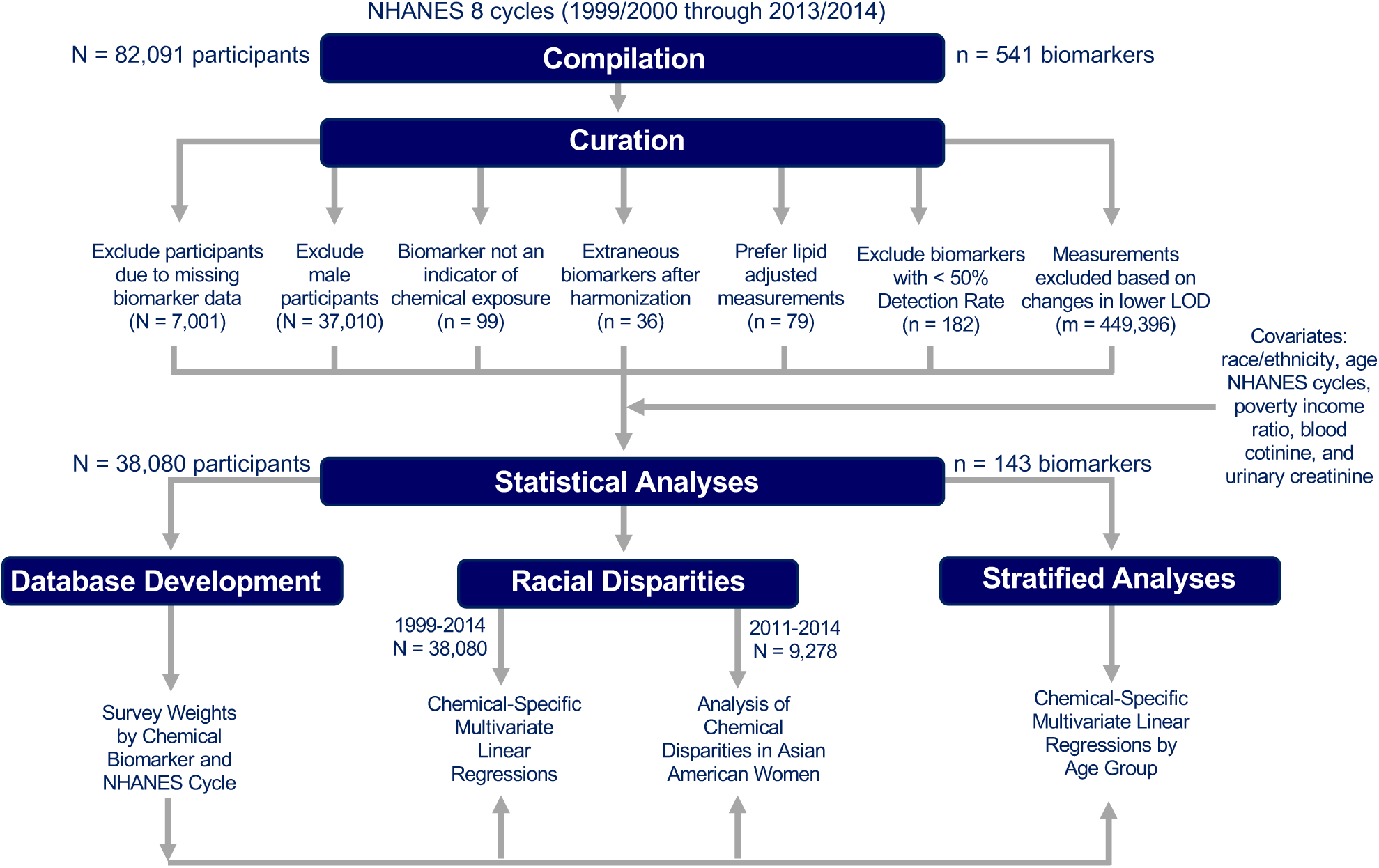
Dataset compilation and cleaning workflow.

### 2.2 Chemical Biomarker Measurements

This section along with **Figure 1** delineate the curation process for selecting chemical biomarkers to include for analysis. First, we excluded biomarkers that are not indicative of chemical exposures (*n* = 99). Next, we corrected for differences in chemical codenames by using a unique codename for each biomarker (*n* = 36). We gave preference to lipid-adjusted data and therefore excluded non-lipid adjusted chemical biomarkers (*n* = 79) when both types of data were provided for a given chemical. We replaced all measurements below the limit of detection (LOD) with 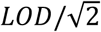 as recommended by the CDC (CDC, 2009). This was to produce reasonably unbiased means and standard deviations (Hornung and Reed, 1990). There were also instances in which urinary cadmium concentrations were recorded as 0 ng/mL due to interference with molybdenum oxide (NCHS, 2005a, NCHS, 2005b). We replaced such values with 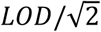 if the participant’s urinary cadmium level was under the LOD or otherwise excluded. We calculated detection frequencies for each chemical biomarker (**Excel Table S2**) and excluded biomarkers with detection frequencies of 50% or less (*n* = 182) across all study participants. Across the NHANES cycles, improvements in laboratory technology can change the LOD and thus lead to differences in detection frequencies by NHANES cycle (Nguyen et al. 2019). To limit bias from these changing LODs over time, we calculated detection frequencies by NHANES cycle (**Excel Table S2**) for each chemical biomarker and excluded measurements that showed drastic changes in the LOD (**Excel Table S3**) and detection frequencies over time (Fig. 1). Measurements from given cycles for several PCBs, Dioxins, Furans, Phytoestrogens, and VOCs along with Paranitrophenol, 2-napthol, 1-pyrene and 9-pyrene (*m* = 449,396) were therefore also excluded based on these criteria (**Excel Table S4**). The final dataset for analysis consisted of 141 chemical biomarkers from 17 different chemical classes (**Excel Table S5**).

### 2.3 Statistical Analysis

We performed all analyses using R version 3.6.0. Given the NHANES complex sampling design, we applied appropriate survey weights in our statistical models to produce estimates representative of the non-institutionalized, civilian US population. To do this, we developed two databases. The first was a database of codenames indicating the appropriate survey weights for each chemical biomarker and NHANES cycle (**Excel Table S6**). For several of the Per- and Polyfluoroalkyl Substance (PFAS), there were two different type of survey weights available within the same cycle (one for children aged 3-11 and the other for participants aged 12 and older). Therefore, we developed another database of codenames indicating which additional survey weights to use when generalizing these results for PFASs (**Excel Table S7**).

Using multivariate regression models, we evaluated differences in biomarker concentrations in blood and urine by race after log-transforming the data. We included log-transformed levels of cotinine as a covariate to represent smoking (Benowitz, 1999), and creatinine levels to adjust for urine dilution and flow differences (Barr et al., 2005). We modeled poverty income ratio (PIR) as a surrogate variable for socioeconomic status. PIR is the ratio of household income and poverty threshold adjusted for family size and inflation. First, we examined the racial differences in chemical biomarker levels by performing a series of chemical-specific regression models with the main predictor being race/ethnicity (categorical), adjusting for age (continuous), sex (categorical), NHANES cycle (continuous), PIR (continuous), and cotinine (continuous) as described in Eq. (1):

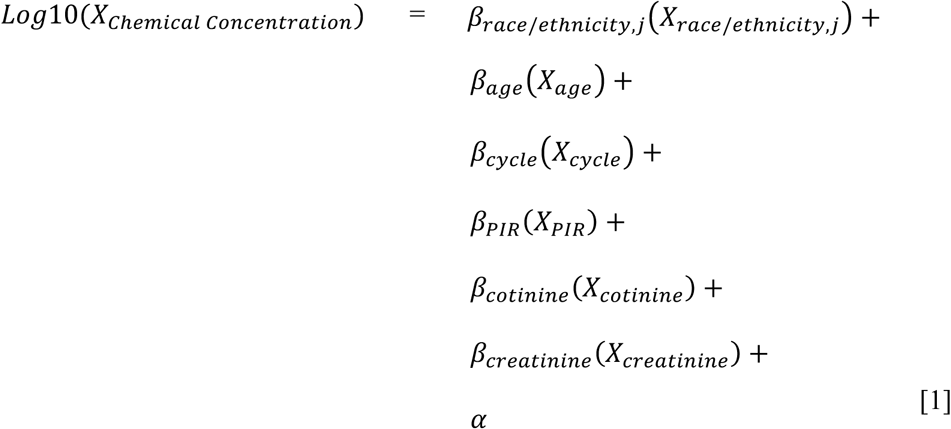

Here, *X*_*Chemical Concentration*_ is the log-transformed, unadjusted chemical biomarker concentration for all participants, *X*_*i*_, where *i* ∈ {*race*/*ethnicity, j, age, sex, cycle, PIR, cotinine, creatinine*}, is the *i* covariate for all participants, *β*_*i*_ is the linear regression coefficient for the *i* covariate, and *α* is the intercept. *X*_*race/ethnicity,j*_, where *j ϵ*{*Mexican Americans, Other Hispanics, Non-Hispanic Black, Other Race*/*Multiracial*} for 1999-2014, is the race covariate for comparing the *jth* race to the reference group of Non-Hispanic Whites. For chemical biomarkers which were measured in urine, we further corrected the regression models by adjusting for urinary creatinine levels (continuous). For the analyses where cotinine concentration was the outcome, the regression models were not further corrected for smoking. Prior to 2011, Asian Americans were categorized in Other Race/Multi-Racial category. Accordingly, to evaluate chemical exposure disparities in Asian American women, we also applied Eq. 1 to the 2011-2014 data. Then to determine whether racial disparities are driven by differences in socioeconomic status, we conducted a sensitivity analysis to observe how the race coefficients change with and without adjustment for PIR in the regression models. The coefficient for *jth* race represents the difference in log-transformed chemical biomarker concentration between the *jth* race and the reference group of Non-Hispanic Whites. To account for multiple comparisons, we used a False Detection Rate (FDR) method on the p-values of the linear regression race-coefficients (Benjamini and Hochberg, 1995).

To evaluate how these racial differences in chemical exposures differ by age group, we conducted stratified analyses by age groups in the 1999-2014 data. We defined 4 age groups: 0-11, 12-25, 26-50, and 51-85. For each age group with chemical biomarker measurements, we performed a chemical specific linear regression with the main predictor as race/ethnicity (categorical) and adjusted for age (continuous), sex (categorical), NHANES cycle (continuous), PIR (continuous), and cotinine (continuous), stratified by age group described in Eq. (2).

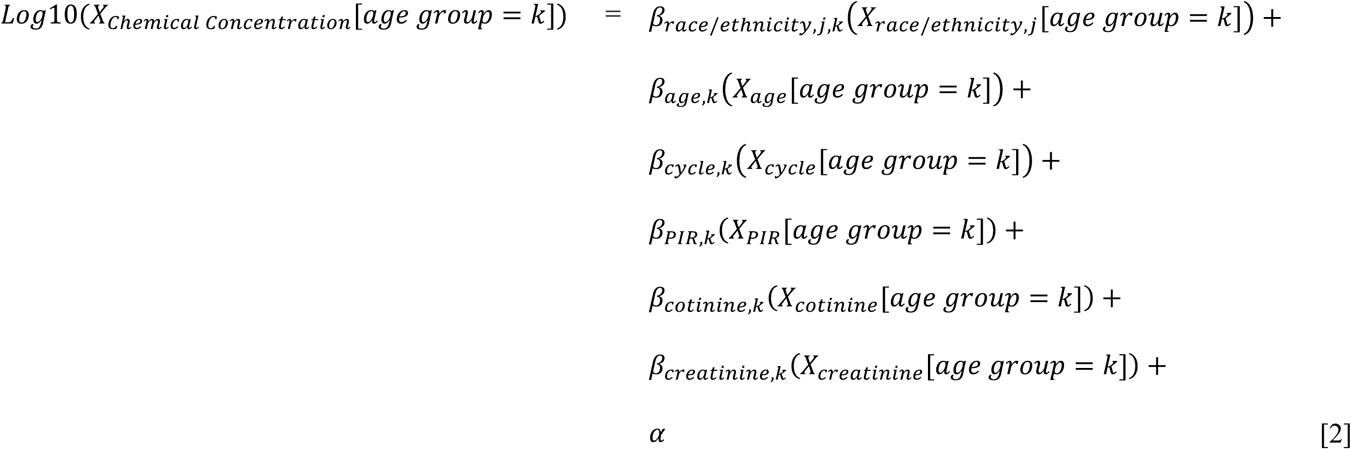

Here, *k* is an available age group from the set of {0-11, 12-25, 26-50, 51-85}, *X*_*Chemical Concentration*_[*age group* = *k*] is the log-transformed, unadjusted chemical biomarker concentration for all participants with ages in the *kth* age groups, *X*_*i,k*_[*age group* = *k*], where *i* ∈ *{race/ethnicity, j, age, sex, cycle, PIR, cotinine, creatinine*}, is the *i* covariate for all participants with ages with the *kth* age group, *β*_*i,k*_ is the linear regression coefficient for the *i* covariate and *kth* age group, and *α* is the intercept. *X*_*race/ethnicity,j,k*_, where *j ϵ*{*Mexican Americans, Other Hispanics, Non-Hispanic Black, Other Race*/*Multiracial*}, is the race covariate for comparing the *jth* race to the reference group of Non-Hispanic Whites in the *kth* age group. To account for multiple comparisons, we used a False Detection Rate (FDR) method on the p-values of the linear regression race-coefficients across all age groups (Benjamini and Hochberg, 1995).

## 3. Results

**Table 1** displays demographic characteristics of the study population. The study population includes 38,080 female study participants of ages 1-85 years, with a median age of 26. Using a series of covariate adjusted regression models, we first calculated the fold-difference in chemical biomarker concentrations by race across the entire study population. These regression results are presented in graphical format in **Figure 2**, where the letters in the plot reflect the fold-difference in chemical biomarkers for each race/ethnicity, relative to non-Hispanic white women, who made up the largest portion of the study population. Full regression results for all covariates in the regression models for each covariate are presented in **Excel Table S8**. Pesticides and pesticide metabolites, including 2,5-dichlorophenol, o,p’-DDE, beta-hexachlorocyclohexane, and 2,4-dichlorophenol had amongst the highest average fold difference across non-Hispanic Black, Mexican American, Other Hispanic, and other race/multiracial women. On average, large differences by race are also apparent for personal care and consumer product compounds including methyl paraben, propyl paraben, monoethyl phthalate and metals, such as mercury and arsenic. Conversely, cotinine, PBDE-153, PBB-153, Equol, DEET, and bisphenol F were among the chemicals of which non-Hispanic white women had the highest levels.

**Table 1.**
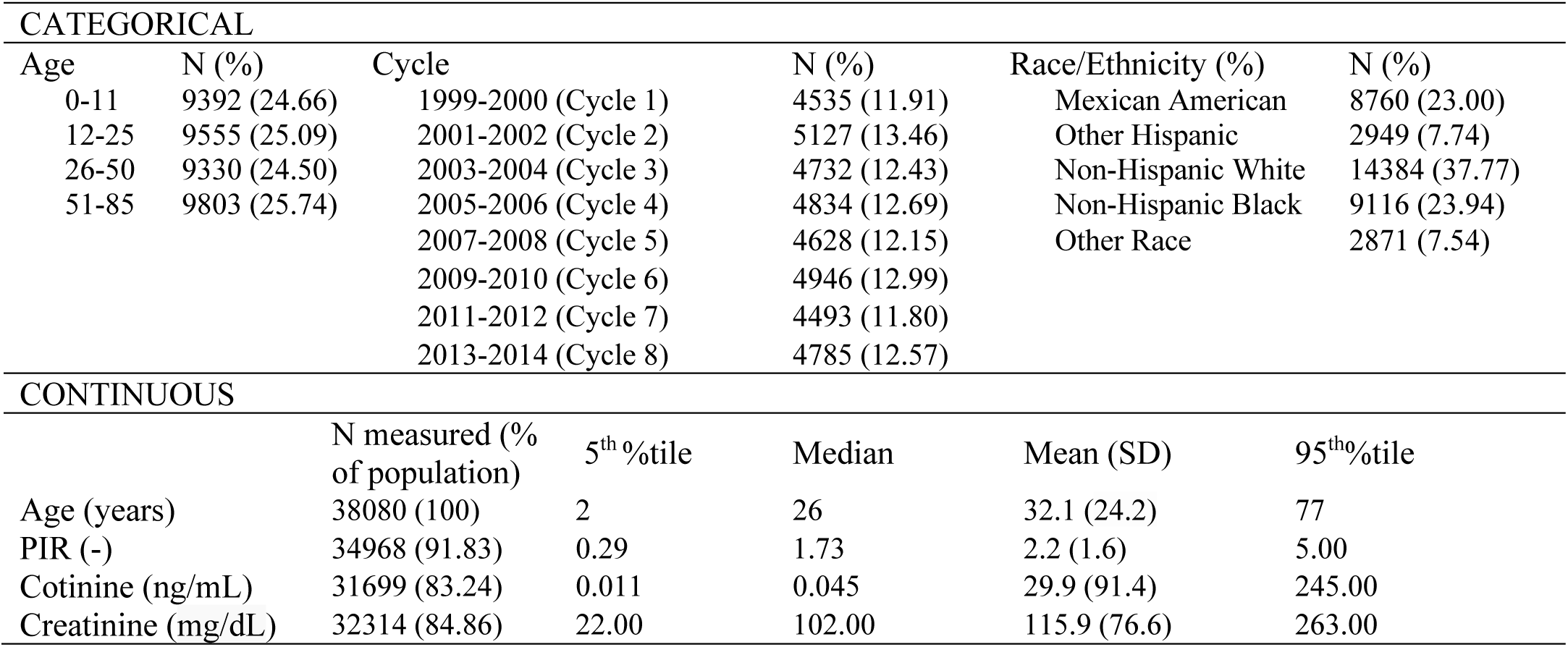
Demographic characteristics of the study population.

**Figure 2.**
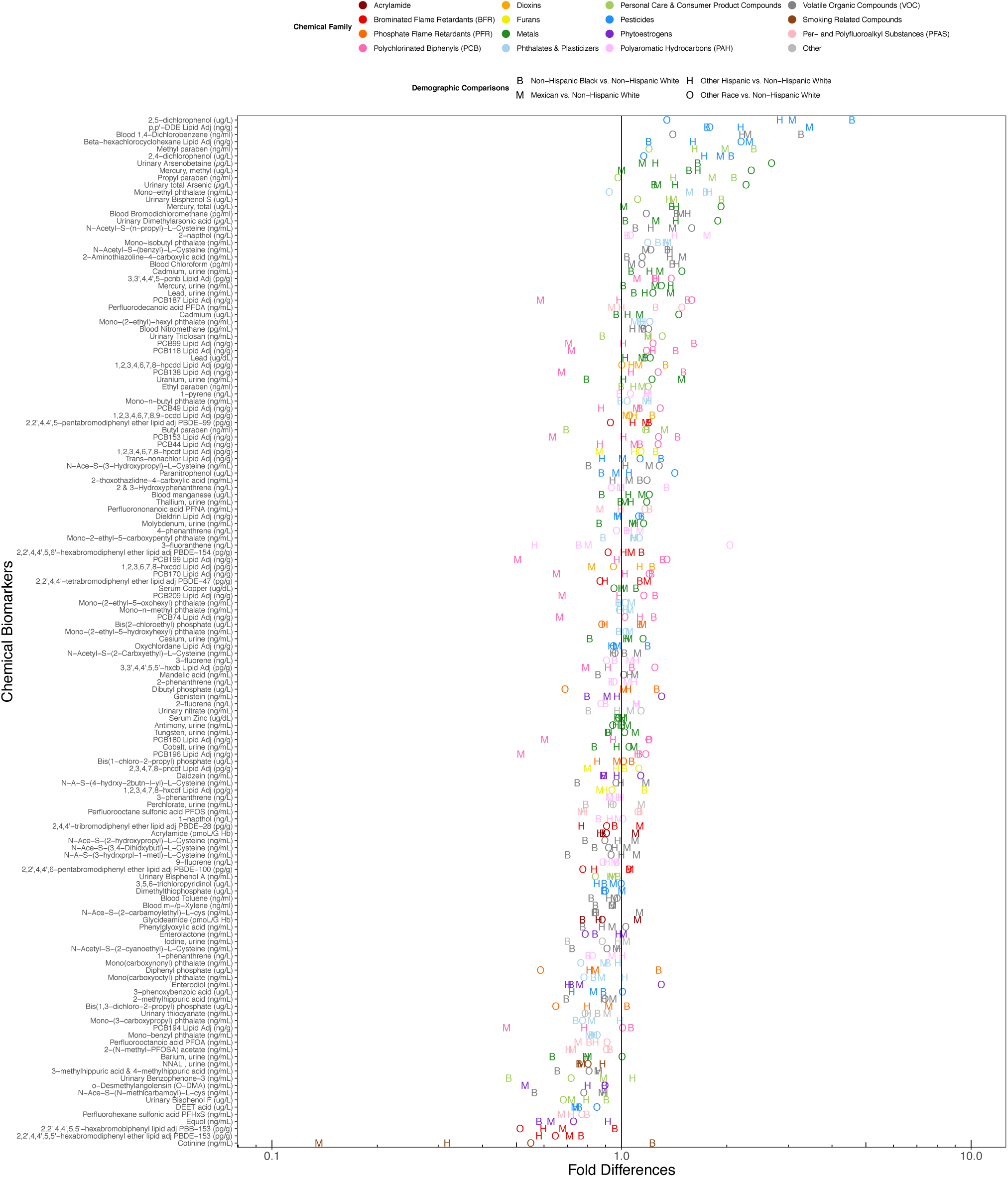
Alphabet soup plot displaying the covariate adjusted fold differences in chemical biomarker concentration by race, ranked by the average difference with non-Hispanic White individuals. Colors represent the chemical families. Shapes represent the comparison between a given race and non-Hispanic White individuals.

In order to more clearly visualize the differences in chemical biomarkers by race/ethnicity, we generated volcano plots, which are displayed in **Figure 3**. The x-axis of these plots depicts the fold difference in average chemical biomarker concentration between each race/ethnicity and non-Hispanic white women. The y-axis depicts statistical significance, as reflected in the negative log_10_ transformation of the FDR-adjusted p-value from the regression analysis for that chemical biomarker, where chemicals with larger values on the y-axis are more statistically significant. As shown in **Figure 3A**, non-Hispanic black women have biomarker concentrations that are more than twice those of non-Hispanic white women for multiple chemicals. These include 2,5-dichlorophenol, 1,4-dichlorobenzene, methyl paraben, monoethyl phthalate, 2,4-dichlorophenol, and propyl paraben. The heavy metals, mercury (p-value = 1.39E-15) and lead (p-value = 1.85E-14), are also significantly higher in non-Hispanic Black women. Conversely, levels of benzophenone-3, a UV blocker used in sunscreen, are significantly higher in non-Hispanic white women (p-value = 1.96E-15). In general, concentrations of PCBs tend to be modestly elevated in non-Hispanic Black women, while volatile organic compounds (VOCs) and phytoestrogen concentrations are higher in non-Hispanic white women. **Figure 3B** shows relative differences in chemical biomarker concentrations between Mexican American and non-Hispanic white women. Pesticides, including 2,5-dichlorophenol, beta-hexachlorocyclohexane, and 2,4-dichlorophenol, along with the polycyclic aromatic hydrocarbon 2-napthol were on average higher in Mexican American women. Conversely, the smoking biomarker, cotinine is significantly lower in Mexican American women (p-value = 8.23E-36). PCB levels, on average, are also lower in Mexican American women, while heavy metal levels tended to be higher. Exposure patterns comparing Other Hispanic and non-Hispanic white women, displayed in **Figure 3C**, showed some similarities, with pesticides 2,5-dichlorophenol and p,p’-DDE elevated in Other Hispanic women. Multiple PFASs, including PFOS, PFHxS, and 2-(N-methyl-PFOSA) acetate, as well as cotinine, are significantly lower in Other Hispanic women. **Figure 3D** shows a distinct exposure pattern in women of other race/ethnicity or multiracial women. Here, levels of heavy metals, including cadmium, mercury, and multiple arsenic biomarkers, are significantly elevated relative to non-Hispanic white women. Conversely, the smoking biomarkers, NNAL (p-value = 2.77E-07) and cotinine (p-value = 4.49E-4), are significantly lower.

**Figure 3.**
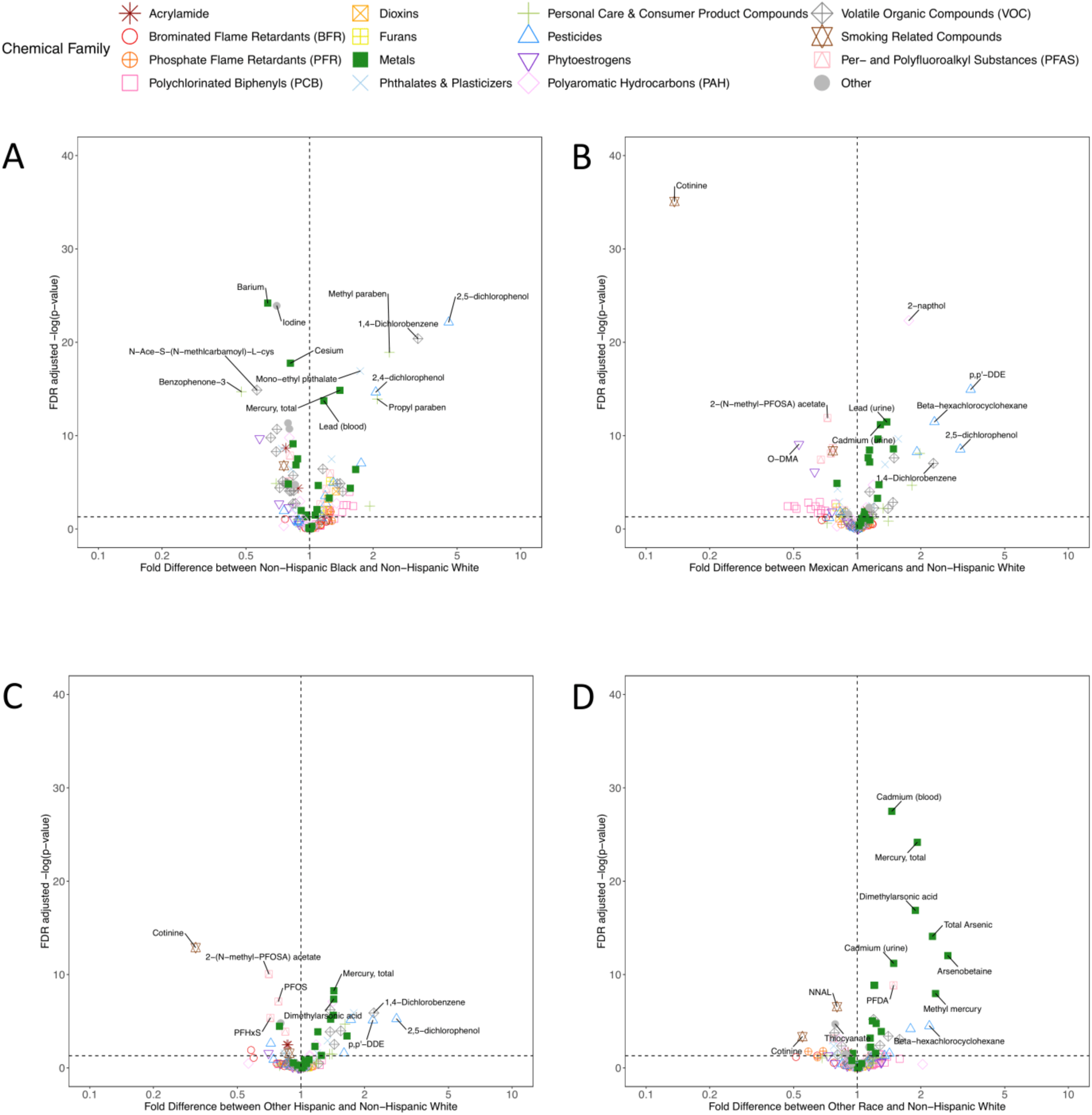
Volcano plots representing the significance of the covariate-adjusted differences in chemical biomarker concentrations between non-Hispanic white women and (A) non-Hispanic Black women, (B) Mexican American women, (C) Other Hispanic women, and (D) Other race/multiracial women. Color and shapes represent the chemical families.

To understand whether socioeconomic status is a driver of racial disparities in chemical exposures, we generated a series of correlation plots, comparing how the differences in chemical biomarker concentrations by race/ethnicity change with the inclusion and exclusion of PIR in the regression models (**Figure S1** and **Excel Table S9**). For many of the chemicals, the fold differences for comparing chemical biomarker levels by race did not change drastically when including PIR as a covariate in the regression models, implying that socioeconomic status is not the primary driver in explaining differences in chemical exposures. However, for cotinine, PCB 194, and several chemicals used in personal care products, the relative differences changed by greater than 25% when PIR was included as a covariate in the regression models. This suggests that either exposure differences between races for these chemicals are mediated by PIR, and/or exposure differences are explained by interactions between race and socioeconomic status. To visualize differences in chemical biomarker concentrations by race across a gradient of income for a few selected biomarkers, we generated violin plots of the chemical biomarker distribution stratified by categories of PIR for each race/ethnicity (**Figure S2**). For benzophenone-3 and cotinine (**Figure S2A and S2B**), the trends of biomarker concentrations across the PIR categories and the average concentrations within the same PIR categories differ by race. This is similar for ethyl paraben (**Figure S2C**), but differences are not as drastic. On the other hand, mercury (**Figure S2D**) along with other remaining chemicals demonstrated a very different pattern from those of the previously mentioned substances. Across all races, the trends across PIR categories are similar for mercury, but within the same PIR category, there are differences in biomarker concentrations by race, suggesting that many chemical exposures disparities by race are independent of PIR.

Starting in 2011, more detailed information on NHANES study participant race/ethnicity were collected, including specifically identifying individuals who report Asian ethnicity. To understand whether the results presented in **Figure 3D** predominantly reflect results in Asian women, who prior to 2011 were categorized in other race/multi-racial category, we assessed exposure disparities specifically in the Asian population. These results, presented in **Figure 4A**, show that, on average, multiple heavy metal biomarkers are more than 2-fold higher relative to non-Hispanic white women, including cadmium, mercury, lead, and arsenics. Additionally, the PFAS compound PFDA is significantly higher in Asian women (p-value = 3.82E-06), while cotinine (p-value = 1.88E-05) and biomarkers of phosphate flame retardants (Bis(1,3-dichloro-2-propyl) phosphate p-value = 5.41E-3; Dibutyl phosphate p-value = 6.76E-4; Diphenyl phosphate p-value = 3.27E-3) are significantly lower. We also calculated whether there were significant disparities in chemical biomarker concentrations in women of other or multi-race after excluding Asian women. **Figure 4B** suggests relatively few differences in this regard, confirming that the other race effect in **Figure 3D** is indeed associated with Asian women. Full regression results across all covariates for the 2011-2014 data are presented in **Excel Table S10**.

**Figure 4.**
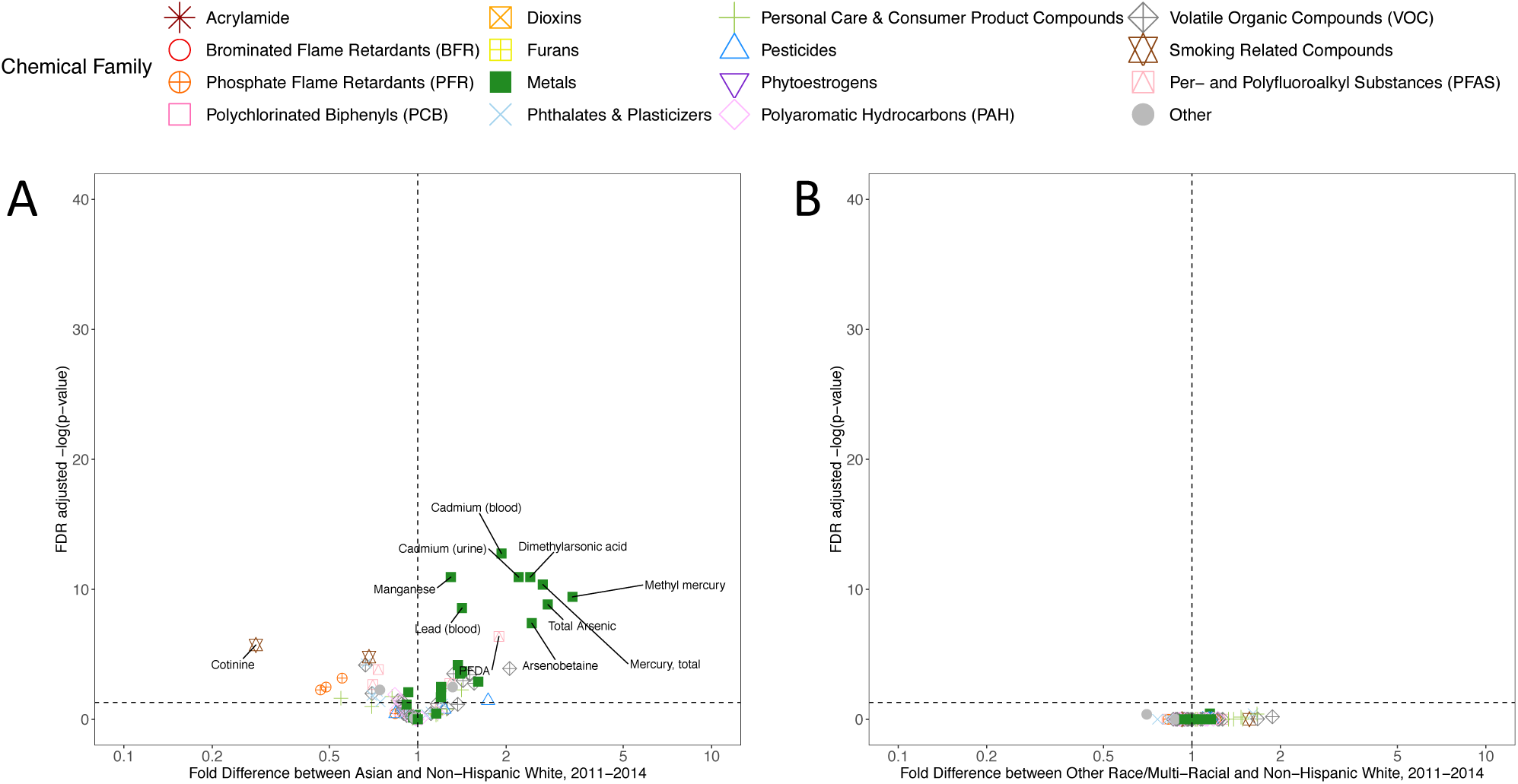
Volcano plots representing the significance of the covariate-adjusted differences in chemical biomarker concentrations between non-Hispanic white women and (A) Asian women, and (B) other race /multiracial women in NHANES 2011-2014. Colors and shapes represent the chemical families.

We have previously shown dramatic differences in the chemical “exposome” by age in NHANES study participants, not stratified by gender or race (Nguyen et al. 2019). Here, we tested for differences in chemical biomarkers by race, after stratifying by age group. **Figure 5** displays these results across the entire study population from 1999-2014. **Excel Tables S11-S14** includes the results for all regression analyses stratified across each of the four age groups. Blue colors reflect chemicals where levels are higher in non-Hispanic white women, while red colors reflect chemicals that are of higher concentration in women of the labeled race/ethnicity. Here, there appear to be exposure disparity patterns that persist across age groups – such as higher 2,4- and 2,5-dichlorophenol concentrations in Mexican American, Other Hispanic, and non-Hispanic black women. Differences in 1,4-dichlorobenzene concentrations are consistent across age groups, although this biomarker was not measured in the youngest individuals. Heavy metal concentrations are elevated in women of other race across age groups. Some exposure patterns differ by age, however. For example, differences in methyl and propyl paraben are most apparent between young non-Hispanic black and non-Hispanic white women less than 12 years old. Increased levels of phosphate flame retardants and the insect repellent DEET in non-Hispanic white women are the most evident in women less than 12 years of age. Similarly, higher relative concentrations of benzophenone-3, bisphenol A, and bisphenol F occur in non-Hispanic white women less than 12. Elevated PCB levels in non-Hispanic black women shown in **Figure 3A** are most evident in women greater than 51 years of age. Overall, these results highlight racial exposure disparities that are either stable or that vary across age groups.

**Figure 5.**
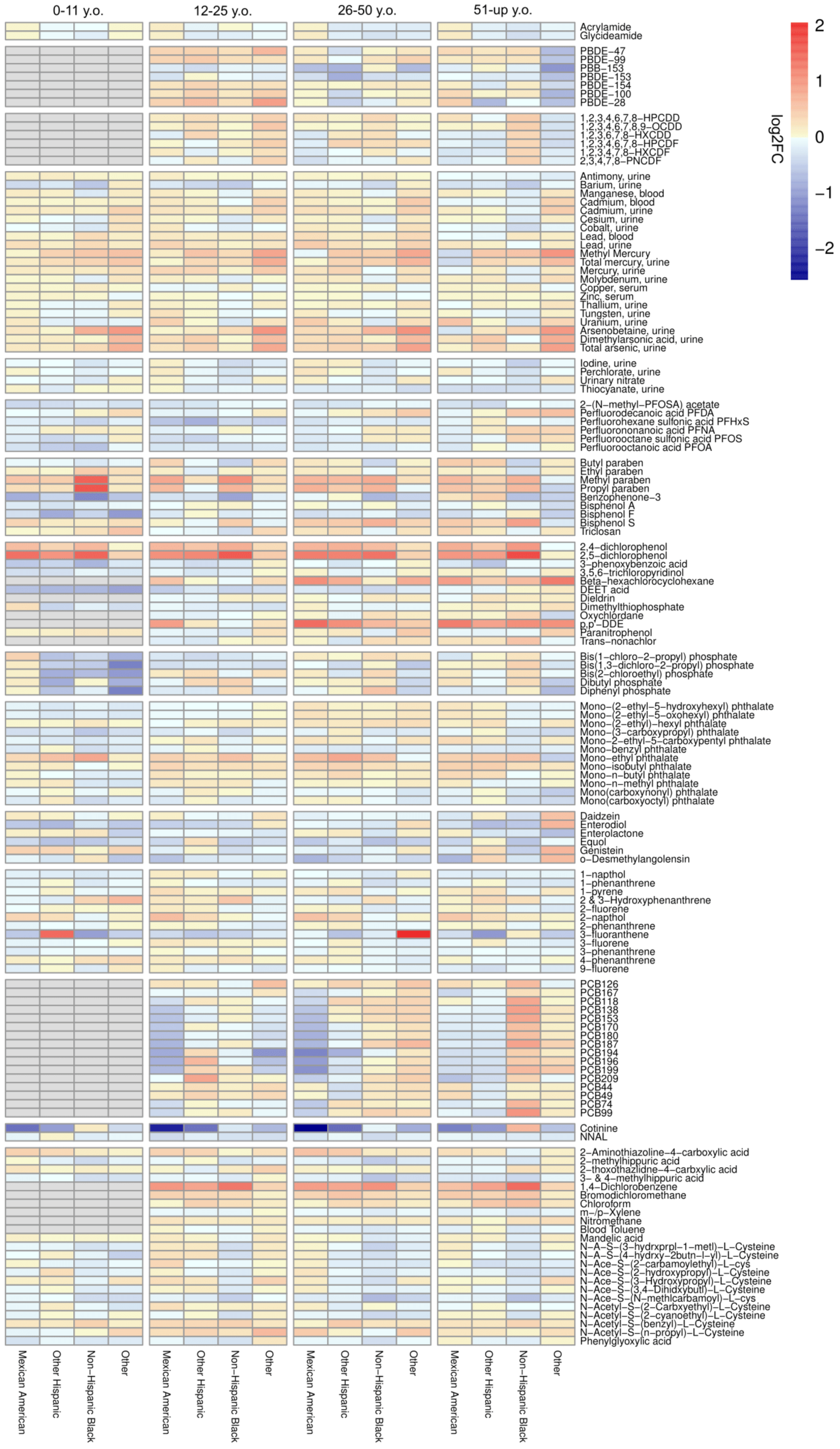
Heatmap displaying covariate adjusted fold differences in chemical biomarker concentrations by race, relative to non-Hispanic white women, stratified by age group and chemical family. Color reflects the log2 fold difference in chemical biomarker concentration. Biomarkers in grey color were not measured in that age group.

## 4. Discussion

Based on population based chemical biomonitoring generated as part of the 1999-2014 NHANES, we performed a comprehensive analysis of racial disparities in biomarker concentrations of 141 chemicals in 38,080 participants. Specifically, we quantified the relative magnitude of racial disparities for individual chemicals and chemical families while utilizing appropriate regression weightings. This helped ensure that the results were as generalizable to the entire US population. These results highlighted striking differences in chemical biomarker exposure patterns by race/ethnicity, independent of other demographic factors such as socioeconomic status. In particular, exposure patterns of pesticides, heavy metals, tobacco smoke associated compounds, and chemicals found in personal care products are found to be most disparate across race/ethnic groups. Stratified analyses revealed exposure patterns that persisted across age groups. For example, this was apparent in heavy metals exposure for women who identify as other race or multiracial, as well as in age-specific exposure patterns, such as elevated PCB, dioxin, and dibenzofuran exposure in older non-Hispanic black women. In some cases, average differences in chemical biomarker concentrations between race/ethnic groups exceeded 400%, such as for urinary propyl or methylparaben concentrations between the youngest non-Hispanic Black and non-Hispanic white women. These findings contextualize racial disparities in chemical exposures across US women and highlight the vast differences in chemical exposomes between demographic groups with well characterized disparities in health outcomes.

Environmental injustice is the disproportionate exposure of individuals of color, lower socioeconomic status, or other politically disadvantaged groups to toxic chemicals in food, air, consumer products, at the workplace, or in their communities (Brulle and Pellow 2005). Disproportionate chemical exposures have been hypothesized to be important drivers of health disparities, including obesity and neurodevelopmental outcomes (Landrigan et al. 2010). While the primary goal of this study was to quantify and compare chemical exposure disparities across racial/ethnic groups, independent of income, others have evaluated combined income and race related disparities in exposure. For instance, one analysis compared geometric mean concentrations of 228 chemical biomarkers between six groups stratified by income and race in NHANES and identified 37 chemicals as likely contributing to environmental justice (Belova et al. 2013). Some of these chemicals, including cotinine, lead, 2,4- and 2,5-dichlorophenol, methyl paraben, and propyl paraben, were associated with the highest disparities across race/ethnic group in the present study. We also compared chemical exposures disparities across racial/ethnic groups with and without adjustment for income and found that cotinine, PCB 194, methyl mercury, and chemicals used in personal care products such as benzophenone-3, the parabens, and triclosan show disparities across both race and socioeconomic status. However, for most of the studied chemicals, differences in chemical exposures were not driven by socioeconomic status but were instead primarily associated with race/ethnicity. Furthermore, a study of racial and social disparities in exposure to BPA and PFAS examined differences in biomarker concentrations in NHANES study participants (Nelson et al. 2012). The concentrations of the four PFAS chemicals examined, PFOA, PFOS, PFNA, and PFHxS, were inversely associated with household income, while BPA concentrations were higher in individuals who reported low food security (Nelson et al. 2012). Here, we identified that, independent of socioeconomic status, as assessed by poverty-income ratio, non-Hispanic white women had the highest concentrations of PFOA, while non-Hispanic Black and other race/multiracial women had the highest concentrations of PFDA. Major routes of exposure to PFAS compounds include contaminated drinking water (Hu et al. 2016), diet (Schecter et al. 2010), and occupational routes (Laitinen et al. 2014). BPA concentrations were not strikingly different by race in our study, but non-Hispanic Black women had, on average, 93% higher BPS concentrations than non-Hispanic white women. Common routes of exposure to BPA and other bisphenol analogues are diet, thermal paper, and personal care products (Chen et al. 2016). Further research is necessary to identify the major routes of exposure which are driving racial disparities in PFAS and bisphenol chemicals biomarker concentrations.

The findings of highly elevated monoethyl phthalate and methyl and propyl paraben concentrations in the non-Hispanic Black women is consistent with a personal care product route of exposure. A study assessing the chemical composition of hair products used by Black women consistently identified high levels of cyclosiloxanes, parabens, and the fragrance carrier diethyl phthalate (Helm et al. 2018). In our study, the concentrations of the diethyl phthalate metabolite monoethyl phthalate were approximately 78% higher on average in non-Hispanic black women of all ages relative to non-Hispanic white women, and 122% higher in non-Hispanic black women less than 12 years of age. This is concerning, since urinary concentrations of monoethyl phthalate have been positively associated with odds of developing breast cancer in a case-control study of women from Northern Mexico (López-Carrillo et al. 2010). Differences in concentrations of methyl and propyl paraben biomarkers were among the highest observed in this study, particularly for the youngest non-Hispanic Black women. These chemicals have been used as preservatives in personal care products, pharmaceuticals, and food additives, and have been found to promote cell growth through multiple mechanisms, including estrogenicity (Gonzalez et al. 2018, 2019; Okubo et al. 2001) and epidermal growth factor receptor signaling (Pan et al. 2016). Particularly relevant to our findings of the greatest methyl and ethyl paraben disparities in the youngest non-Hispanic Black women was the finding that early life paraben exposures can alter developing mammary gland morphology and induce gene expression that resembles an early cancer-like state (Gopalakrishnan et al. 2017). Use of hair products has been identified as a potential risk factor for breast cancer in non-Hispanic Black women (Stiel et al. 2016). Further research is needed, however, to determine whether early-life exposure to potentially estrogenic compounds, like parabens, can induce biological alterations that increase risk of estrogen receptor negative breast cancers.

One of the most apparent disparities in chemical biomarker concentrations by race was with the compounds 2,4-dichlorophenol, 2,5-dichlorophenol, and 1,4-dichlorobenzene. 1,4-dichlorobenzene is used as a disinfectant, pesticide, and deodorant. 2,5-dichlorophenol is a metabolite of 1,4-dichlorobenzene, while 2,4-dichlorophenol is a metabolite of the antimicrobial triclosan or other pesticides. Elevated concentrations of these chemicals in non-Hispanic Black individuals has been noted previously (Belova et al. 2013; Ye et al. 2014) The concentrations of these three chemicals were up to 350% higher on average in non-Hispanic Black women, relative to non-Hispanic white women, and also elevated in Mexican American and Other Hispanic women. Importantly, these exposure disparities were consistent across all age groups. While 2,4-dicholorophenol concentrations were significantly elevated in non-Hispanic Black and Hispanic women, urinary triclosan levels were not significantly different by race/ethnicity. This suggests that either triclosan is not the main chemical exposure that explains the differences in concentrations of 2,4-dichlorophenol or that there are differences in metabolism and excretion rates by race, which is less likely. 1,4-dichlorobenzene exposure has been associated with altered thyroid biomarkers in NHANES (Wei and Zhu 2016), altered immunologic and liver function parameters in occupationally exposed workers (Hsiao et al. 2009), and altered sperm production and increased prostate weight in exposed rats (Takahashi et al. 2011). Understanding and mitigating exposure to these chemicals is therefore of importance to reduce disparate risk of these health outcomes.

Heavy metals were among the chemicals most consistently different across racial/ethnic groups. In particular, women who identified as other race or multiracial had the highest concentrations of multiple metals, including cadmium, mercury, arsenics, lead, and manganese. Focusing on data from NHANES 2011-14, we identified that these elevated metals concentrations were restricted to women who identified as Asian. This is consistent with a previous finding of increased concentrations of a subset of these metals in Asian NHANES participants (Awata et al. 2017). Furthermore, elevated levels of mercury, lead, and arsenics were also identified in non-Hispanic Black women, relative to non-Hispanic white women. Mexican American women had elevated levels of uranium, lead, mercury, arsenics, and cadmium, while Other Hispanic women had higher concentrations of mercury, arsenics, and cadmium than non-Hispanic white women. Non-Hispanic white women, however, had higher concentrations of urinary barium. Previous research has linked diet, occupation, education level, and smoking status to elevated metals exposure (Awata et al. 2017), in addition to housing (Jacobs et al. 2013), air pollution (Suvarapu and Baek 2016), and contaminated water (Pieper et al. 2017). The well characterized toxicity of heavy metals exposure, even at low doses, make identifying and ameliorating heavy metal exposures a top priority for addressing environmental health disparities.

The oldest non-Hispanic Black women in our study had consistently higher concentrations of persistent organic pollutants, including dioxins, dibenzofurans, PCBs, and DDT metabolites. This is consistent with a previous report of non-Hispanic black individuals having an increased risk of having multiple persistent organic pollutants detectable their blood (Pumarega et al. 2016) or higher average levels of PCBs (Xue et al. 2014). Biomarkers of persistent organic pollutants were quantified on an individual (non-pooled) basis in the 1999-2004 NHANES cycles. Elevated concentrations of these pollutants, such as the DDT metabolite, DDE, have been associated with an increased risk of breast cancer (Wolff et al. 1993). A lack of disparities, and decreasing concentrations of these chemicals in younger individuals over time, generally reflect a public health success in decreasing population exposures to these toxic compounds (Nguyen et al. 2019). The long half-life of these chemicals suggests that the detected biomarkers predominantly reflect historical exposures. This could, however, be of substantial importance for children of non-Hispanic Black women, who could have been exposed to disproportionately high levels of these chemicals in the womb or early in childhood. For example, *in utero* exposure to the pesticide, DDT, has been associated with an increased risk of breast cancer in adulthood. Specifically, women in the highest quartile of *in utero* DDT exposure were found to have a 3.7-fold increased risk of developing breast cancer relative to women in the lowest quartile of exposure (Cohn et al. 2015). Prenatal exposure to organochlorine compounds has also been associated with decreased lung function later in life (Hansen et al. 2016), risk of infection in childhood (Dewailly et al. 2000), attention deficit hyperactivity disorder (Sagiv et al. 2010), and obesity (Mendez et al. 2011). If these effects of elevated early life persistent organic pollutant exposure last throughout the life course, there could be continued adverse health consequences that manifest in those exposed for the foreseeable future.

Our study had important limitations. First, the cross-sectional nature of NHANES allows only a single biomarker measurement per individual. Moreover, since the half-lives of the biomarkers assessed in this study are highly variable (Nguyen et al. 2019), the precision of estimates of long-term exposure largely varies across chemical family. Additionally, this study was not able to assess geographic variation in exposure. Others have identified that persistent organic pollutant exposures in the NHANES cohort varies geographically, with higher DDT metabolite concentrations in individuals residing in the West, and elevated PCB concentrations in individuals residing in the Northeast (Wattigney et al. 2015). Future work is needed to precisely characterize exposure “hot spots,” in order to design intervention studies to reduce exposure disparities. Our study also focused on identifying average differences in biomarker concentrations. By ignoring the extremes of these distributions, we have likely not considered individuals at greatest risk of developing adverse health outcomes. Similarly, our analyses were limited by low detection rates, with 182 chemicals not meeting our inclusion threshold of at least 50% detection in the study population. A more in-depth analysis of differences in detection frequency by race/ethnicity could identify additional chemicals with significant racial disparities. For chemical biomarkers measured in urine, variations in the concentration of urinary creatinine, used as a correction factor for urine dilution, potentially confounds our comparison of exposures between individuals of different races. This is because increased average concentrations of urinary creatinine have been identified for non-Hispanic Black individuals, relative to Mexican American and non-Hispanic white individuals (Barr et al. 2005). While we adjusted for urinary creatinine as a covariate in our regression models, the still may be residual confounding. The large number of chemicals assessed also precluded an in-depth characterization of the various routes of exposure of individual chemicals – this is undoubtedly an essential future direction of research to develop strategies to eliminate exposure disparities. Finally, while we performed analyzed all chemical biomarkers available from NHANES 1999-2014, these chemicals only represent a small proportion of the over 80,000 chemicals estimated to be used in commerce in the United States. Future studies could benefit from an unbiased metabolomics approach to identity disparities in chemical exposures which are not captured in NHANES.

The persistent health disparities between women of different races/ethnicities makes understanding the etiological drivers of these disparities a pressing public health issue. A recent commentary highlighted a lack of knowledge regarding the molecular underpinnings of health disparities. It described how the vast majority of genome sequencing data had been generated in populations of European ancestry (Sirugo et al. 2019). Environmental exposures, however, are hypothesized to be the major driving risk factors for a vast suite of complex diseases (Rappaport and Smith 2010). Even when genetic data has been generated in an equitable fashion, understanding gene-environment interactions and complex disease phenotypes will still require in-depth quantification of environmental exposures. In this study, we have comprehensively identified differences in biomarker of chemical exposure across women of various race/ethnic groups and across age groups. These findings can guide future efforts to understand chemical impacts on health disparities by helping to prioritize chemicals for assessment in epidemiological studies. Additionally, chemicals as identified as highly disparate here can be further prioritized for toxicological assessment relevant to disease outcomes of interest. Finally, these findings can inform public health interventions designed to reduce chemical disparities and promote health equity across the population.

## Acknowledgements

This work was supported by the Ravitz Family Foundation, the Forbes Institute for Cancer Discovery at the University of Michigan Rogel Cancer Center, as well as grants R01 ES028802 and P30 ES017885 from the National Institute of Environmental Health Sciences and grant T32GM070499 of the National Institute of General Medical Sciences.

## Competing Interests Financial Declaration

The authors have no competing financial interests to declare.

