## Supplemental Figures for "A comprehensive analysis of racial disparities in chemical biomarker concentrations in United States women, 1999-2014"

Affiliations: ^1^ Department of Computational Medicine and Bioinformatics, Medical School, University of Michigan, Ann Arbor, MI, USA; ^2^ Department of Environmental Health Sciences, School of Public Health, University of Michigan, Ann Arbor, MI, USA; ^3^ Center for Computational Medicine and Bioinformatics, University of Michigan, Ann Arbor, MI, USA; ^4^ Department of Nutritional Sciences, School of Public Health, University of Michigan, Ann Arbor, MI, USA; ^5^ Rogel Cancer Center, University of Michigan, Ann Arbor, MI, USA

To whom correspondence should be addressed:

Justin A. Colacino, PhD, MPH, MA

Department of Environmental Health Sciences

University of Michigan School of Public Health,

1415 Washington Heights, 6651 SPH I, Ann Arbor, MI, 48109-2029

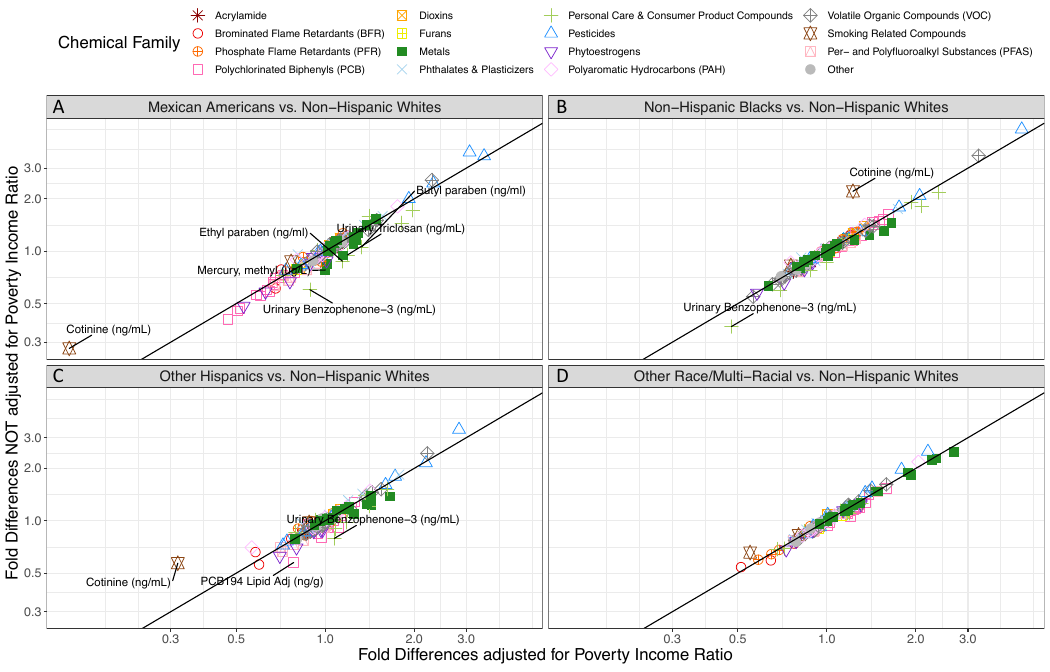

**Figure S1.** Panel of correlation plots comparing fold differences for race that adjusted for poverty income ratio (PIR) with those that excluded PIR from the regression models. Colors and shapes represent the different chemical families. Chemicals were labeled if fold differences changed by greater than 25% when PIR was considered a covariate in the regression models.

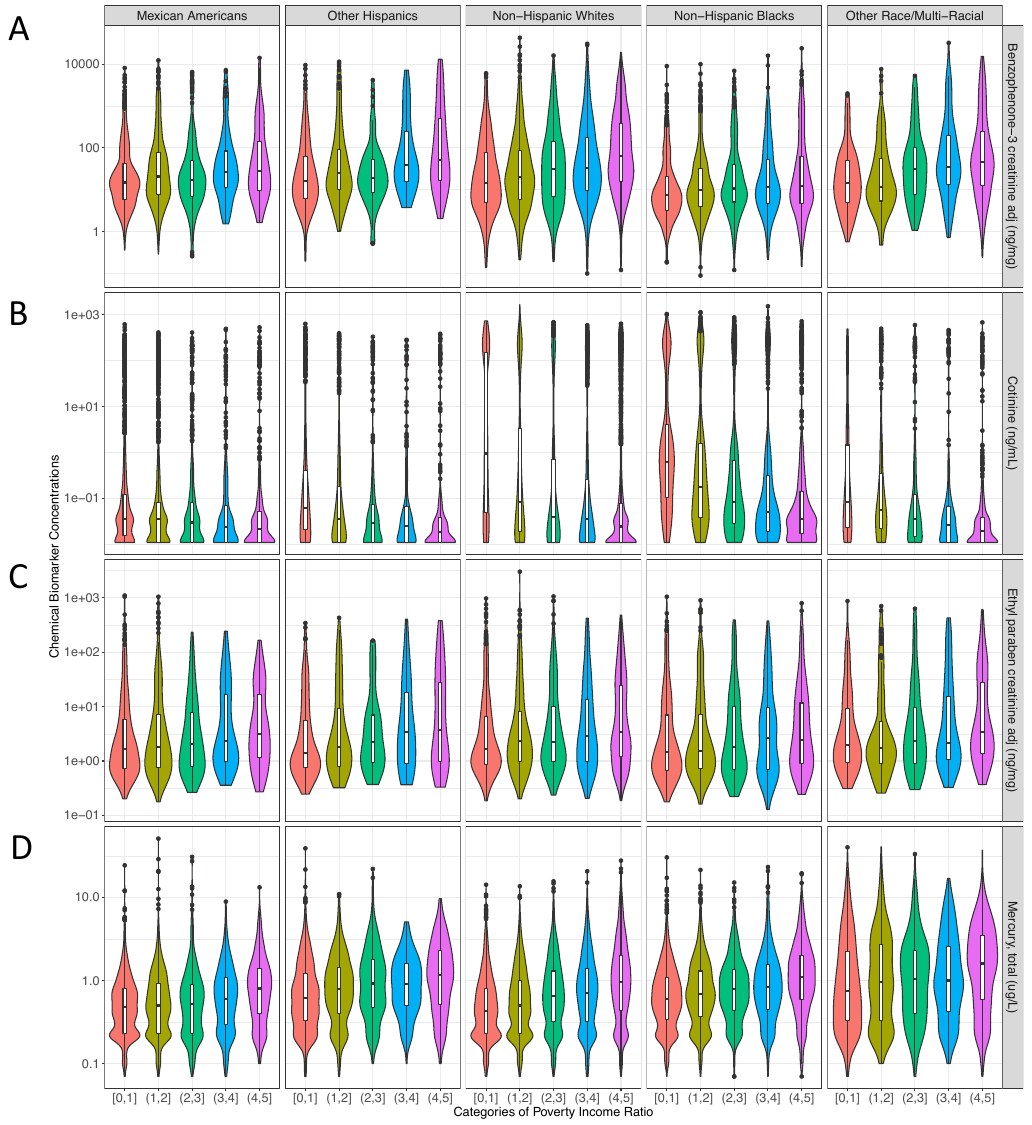

**Figure S2.** Panel of violin plots showing the distribution of chemical biomarker levels changes across categories of poverty income ratio (PIR) and stratified by race for A) an indicator of sunscreen use, benzophenone-3, B) a biomarker of smoking, cotinine, C) a chemical used in personal care products, ethyl paraben, and D) methyl mercury. Colors represent different categories of PIR.
